# ClipperQTL: ultrafast and powerful eGene identification method

**DOI:** 10.1101/2023.08.28.555191

**Authors:** Heather J. Zhou, Xinzhou Ge, Jingyi Jessica Li

## Abstract

A central task in expression quantitative trait locus (eQTL) analysis is to identify cis-eGenes (henceforth “eGenes”), i.e., genes whose expression levels are regulated by at least one local genetic variant. Among the existing eGene identification methods, FastQTL is considered the gold standard but is computationally expensive as it requires thousands of permutations for each gene. Alternative methods such as eigenMT and TreeQTL have lower power than FastQTL. In this work, we propose ClipperQTL, which reduces the number of permutations needed from thousands to 20 for data sets with large sample sizes (*>* 450) by using the contrastive strategy developed in Clipper; for data sets with smaller sample sizes, it uses the same permutation-based approach as FastQTL. We show that ClipperQTL performs as well as FastQTL and runs about 500 times faster if the contrastive strategy is used and 50 times faster if the conventional permutation-based approach is used. The R package ClipperQTL is available at https://github.com/heatherjzhou/ClipperQTL.

## 1 Introduction

Molecular quantitative trait locus (molecular QTL, henceforth “QTL”) analysis investigates the relationship between genetic variants and molecular traits, potentially explaining findings in genome-wide association studies [1, 2]. Based on the type of molecular phenotype studied, QTL analyses can be categorized into gene expression QTL (eQTL) analyses [3, 4], alternative splicing QTL (sQTL) analyses [4], three prime untranslated region alternative polyadenylation QTL (3^*′*^aQTL) analyses [5], and so on [1, 2]. Among these categories, eQTL analyses, which investigate the association between genetic variants and gene expression levels, are the most common. Therefore, in this work, we focus on eQTL analyses as an example, although everything discussed in this work is applicable to other types of QTL analyses as well.

A central task in eQTL analysis is to identify cis-eGenes (henceforth “eGenes”), i.e., genes whose expression levels are regulated by at least one local genetic variant. This presents a multiple-testing challenge as not only are there many candidate genes, each gene can have up to tens of thousands of local genetic variants, and the local genetic variants are often in linkage disequilibrium (i.e., associated) with one another.

Existing eGene identification methods include FastQTL [6], eigenMT [7], and TreeQTL [8]. All three methods share the same two-step approach: first, obtain a gene-level *p*-value for each gene; second, apply a false discovery rate (FDR) control method on the gene-level *p*-values to call eGenes. The key difference between the three methods lies in how the gene-level *p*-values are obtained.

Among the existing eGene identification methods, FastQTL [6] is considered the gold standard and is currently the most popular. It uses permutations to obtain gene-level *p*-values. There are four main ways to use FastQTL, depending on (1) whether the direct or the adaptive permutation scheme is used and (2) whether proportions or beta approximation is used (Table 1). The default way of using FastQTL is to use the adaptive permutation scheme with beta approximation [4, 6]. The adaptive permutation scheme means the number of permutations is chosen adaptively for each gene (between 1000 and 10,000 by default [4, 6]); the beta approximation helps produce higher-resolution gene-level *p*-values given the numbers of permutations (Algorithm S1). The main drawback of FastQTL is the lack of computational efficiency as it requires thousands of permutations for each gene. A faster implementation of FastQTL named tensorQTL has been developed [9], but it relies on graphics processing units (GPUs), which are not universally available.

**Table 1:** Summary of the 11 eGene identification methods we compare. Details of these methods can be found in Sections 4.2 and S1.

|  | Method category | Method | Note | Method name for speed comparison |
| --- | --- | --- | --- | --- |
|  | (A) | (B) | (C) | (D) |
| 1 | Matrix eQTL | Matrix eQTL |  |  |
| 2 | FastQTL | FastQTL_1K-10K_prop |  | FastQTL_1K-10K |
| 3 |  | FastQTL_1K-10K_beta | Default FastQTL method |  |
| 4 |  | FastQTL_1K_prop |  | FastQTL_1K |
| 5 |  | FastQTL_1K_beta |  |  |
| 6 | eigenMT | eigenMT |  | eigenMT |
| 7 | TreeQTL | TreeQTL_BY | Default TreeQTL method | TreeQTL |
| 8 |  | TreeQTL_Storey |  |  |
| 9 | ClipperQTL | ClipperQTL_standard_1K |  | ClipperQTL_standard_1K |
| 10 |  | ClipperQTL_Clipper_20 |  | ClipperQTL_Clipper_20 |
| 11 |  | ClipperQTL_Clipper_50 |  | ClipperQTL_Clipper_50 |

eigenMT [7] and TreeQTL [8] have been proposed as faster alternatives to FastQTL. Neither method uses permutations. In a nutshell, eigenMT uses Bonferroni correction to calculate a gene-level *p*-value for each gene but estimates the *effective* number of local genetic variants for each gene by performing a principal component analysis (conceptually speaking; instead of using the *actual* number of local genetic variants). On the other hand, TreeQTL uses Simes’ rule [10] to calculate a gene-level *p*-value for each gene. Our analysis shows that both eigenMT and TreeQTL have lower power than FastQTL (Figures 1 and 3).

**Figure 1:**
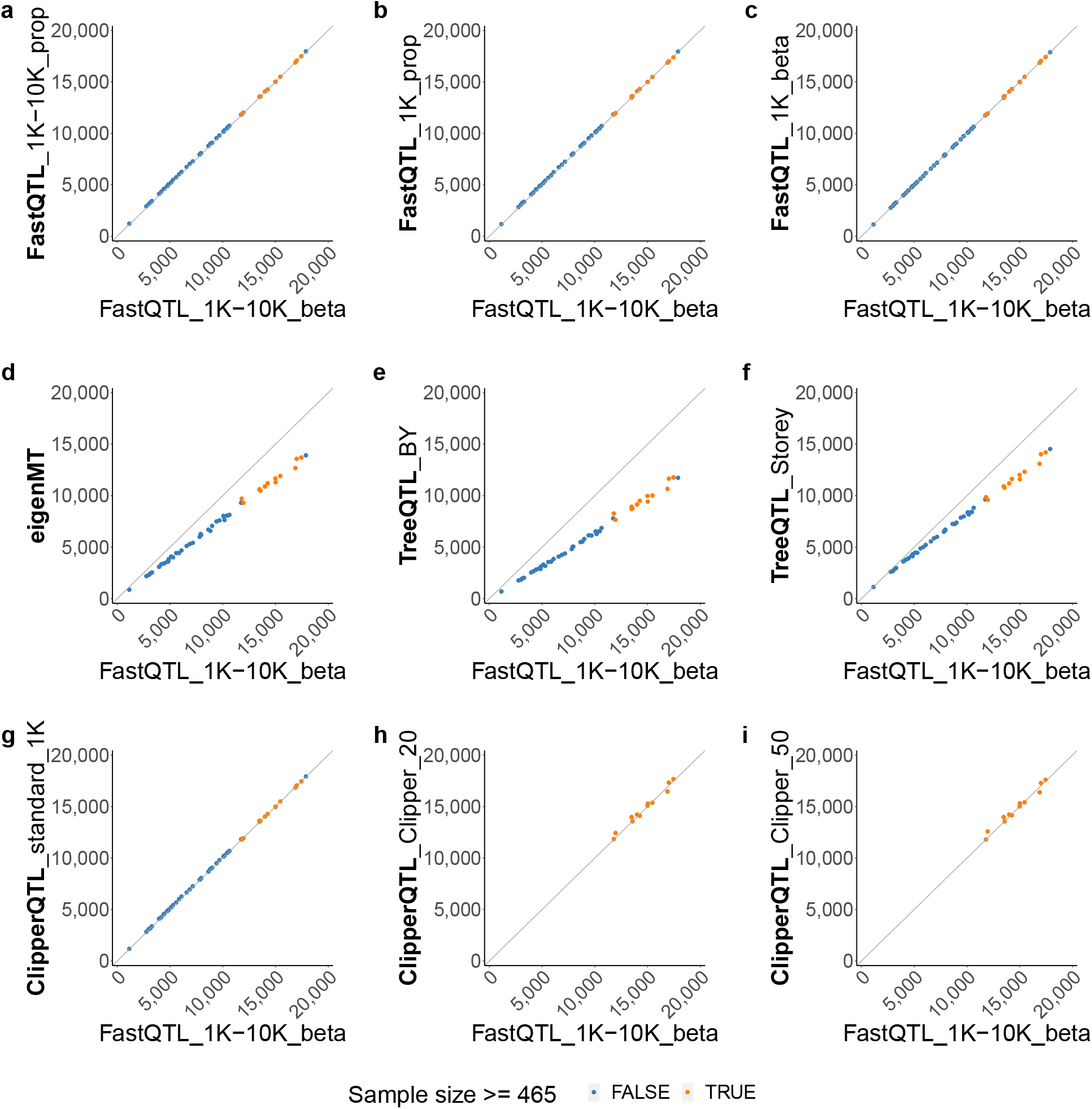
Number of eGenes comparison based on GTEx expression data [4] (Table 1; see Section 2.1 for the analysis details). Each dot corresponds to a tissue. The *x*-axis and *y*-axis both represent numbers of eGenes identified by different methods. Diagonal lines through the origin are shown to help with visualization. **a**-**c** The four variants of FastQTL identify almost the same numbers of eGenes as one another. **d**-**f** eigenMT and TreeQTL methods identify fewer eGenes than. **g**-**i** ClipperQTL methods identify almost the same numbers of eGenes as FastQTL in tissues with the appropriate sample sizes (Section 4.2). We use 465 as the sample size cutoff because the next largest sample size is 396. See Figure S2 for an analysis of the overlap between identified eGenes.

Clipper [11] is a *p*-value-free FDR control method. Given a large number of features (e.g., genes), a number of measurements under the experimental (e.g., treatment) condition, and a number of measurements under the background (e.g., control) condition, Clipper works as the following: first, obtain a contrast score for each feature based on the experimental and background measurements (for example, the contrast score may be the average experimental measurement minus the average background measurement); second, given a target FDR (e.g., 0.05), obtain a cutoff for the contrast scores; lastly, call the features with contrast scores above the cutoff as discoveries. The idea is that the contrast scores of the uninteresting features (e.g., genes whose expected expression levels are not increased by the treatment) will be roughly symmetrically distributed around zero, and the outlying contrast scores in the right tail likely belong to interesting features. Notably, Clipper produces a *q*-value for each feature (similar to Storey’s *q*-values [12]), so that the features can be ranked from the most significant to the least significant.

In this work, we propose ClipperQTL for eGene identification, which reduces the number of permutations needed from thousands to 20 for data sets with large sample sizes (*>* 450) by using the contrastive strategy developed in Clipper; for data sets with smaller sample sizes, it uses the same permutation-based approach as FastQTL. Unlike tensorQTL, our ClipperQTL software does not rely on GPUs. We show that ClipperQTL performs as well as FastQTL and runs about 500 times faster if the contrastive strategy is used and 50 times faster if the conventional permutation-based approach is used (we refer to the two variants of ClipperQTL as the Clipper variant and the standard variant, respectively; Section 4.2).

## 2 Results

### 2.1 Real data results

We compare the performance and run time of different variants of FastQTL, eigenMT, TreeQTL, and ClipperQTL (Table 1) on the most recent GTEx expression data [4]. The 49 tissues with sample sizes above 70 are considered [4]. For each gene, we consider single nucleotide polymorphisms (SNPs) within one megabase (Mb) of the transcription start site (TSS) of the gene [4]; we use 0.01 as the threshold for the minor allele frequency (MAF) of a SNP and 10 as the threshold for the number of samples with at least one copy of the minor allele (MA samples) [6]. We include eight known covariates and a number of top expression PCs (principal components) as inferred covariates [13]. The eight known covariates are the top five genotype PCs, WGS sequencing platform (HiSeq 2000 or HiSeq X), WGS library construction protocol (PCR-based or PCR-free), and donor sex [4]. The number of expression PCs is chosen via the Buja and Eyuboglu (BE) algorithm [13, 14] for each tissue. We use the BE algorithm because we find that in our simulated data (Section S2), the BE algorithm can recover the true number of covariates well. The target FDR for eGene identification is set at 0.05. We do not include Matrix eQTL [15] in our real data comparison because both our simulation study (Section 2.2) and Huang et al. [16] show that Matrix eQTL cannot control the FDR in the eGene identification problem.

The results from our real data analysis are summarized in Figures 1, 2, and S2. We find that the four variants of FastQTL produce almost identical results as one another. Specifically, the numbers of eGenes identified by the four methods are almost identical (Figure 1), and the identified eGenes highly overlap (Figure S2). This means the adaptive permutation scheme and the beta approximation of FastQTL (Section S1.2) are not critical to the performance of FastQTL; the simplest variant, FastQTL_1K_prop, is sufficient. Further, we find that eigenMT and TreeQTL methods identify fewer eGenes than FastQTL (Figure 1). In contrast, ClipperQTL methods produce almost identical results as FastQTL in tissues with the appropriate sample sizes (Section 4.2; Figures 1 and S2).

**Figure 2:**
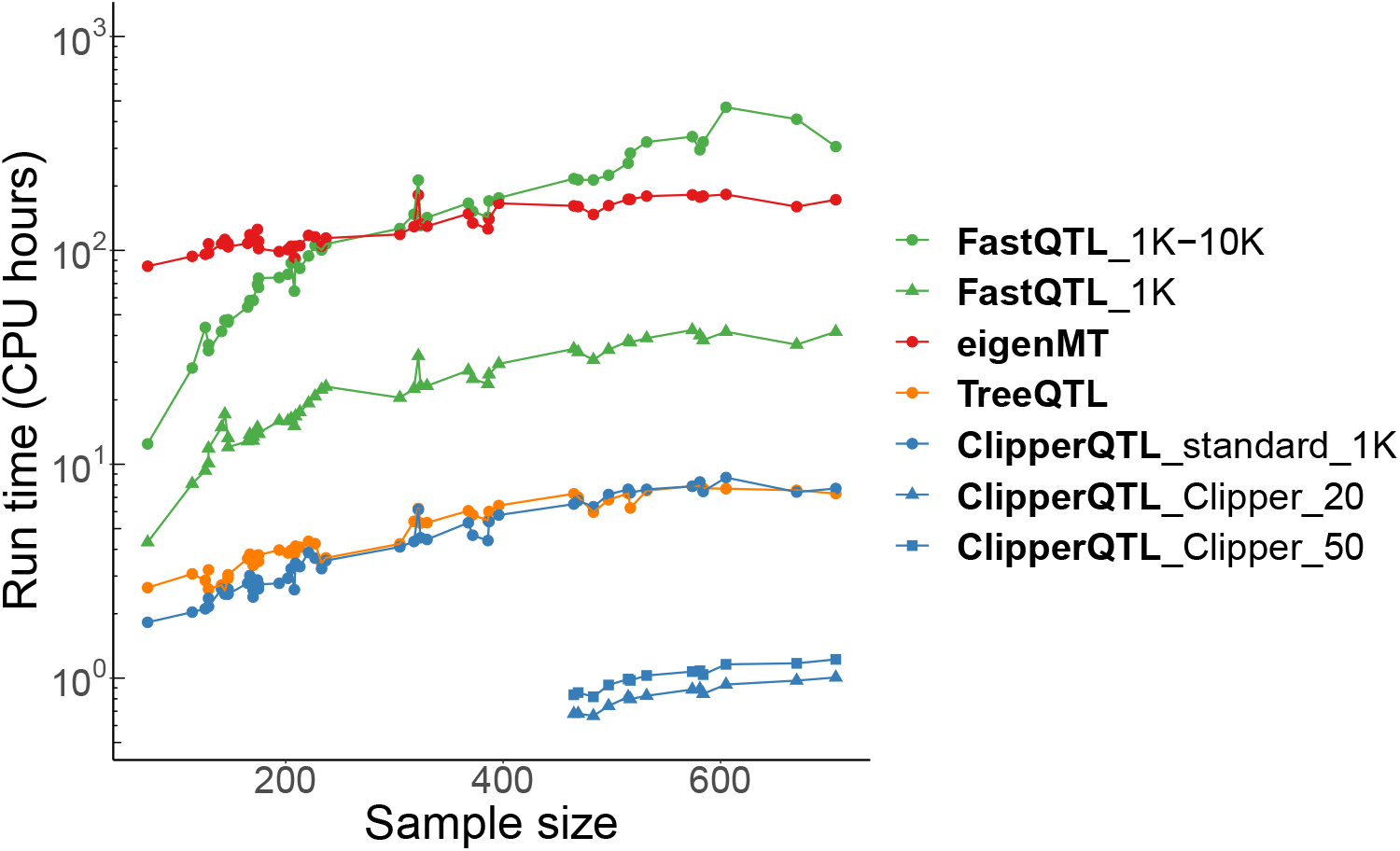
Run time comparison based on GTEx expression data [4] (Table 1; see Section 2.1 for the analysis details). Each dot corresponds to a tissue. FastQTL_1K-10K takes under 500 CPU hours. FastQTL_1K takes under 50 CPU hours. ClipperQTL_standard_1K takes under 10 CPU hours. ClipperQTL_Clipper_20 takes under 1 CPU hour. Run times of ClipperQTL_Clipper_20 and ClipperQTL_Clipper_50 are only shown for tissues with sample sizes ≥ 465 (Figure 1).

**Figure 3:**
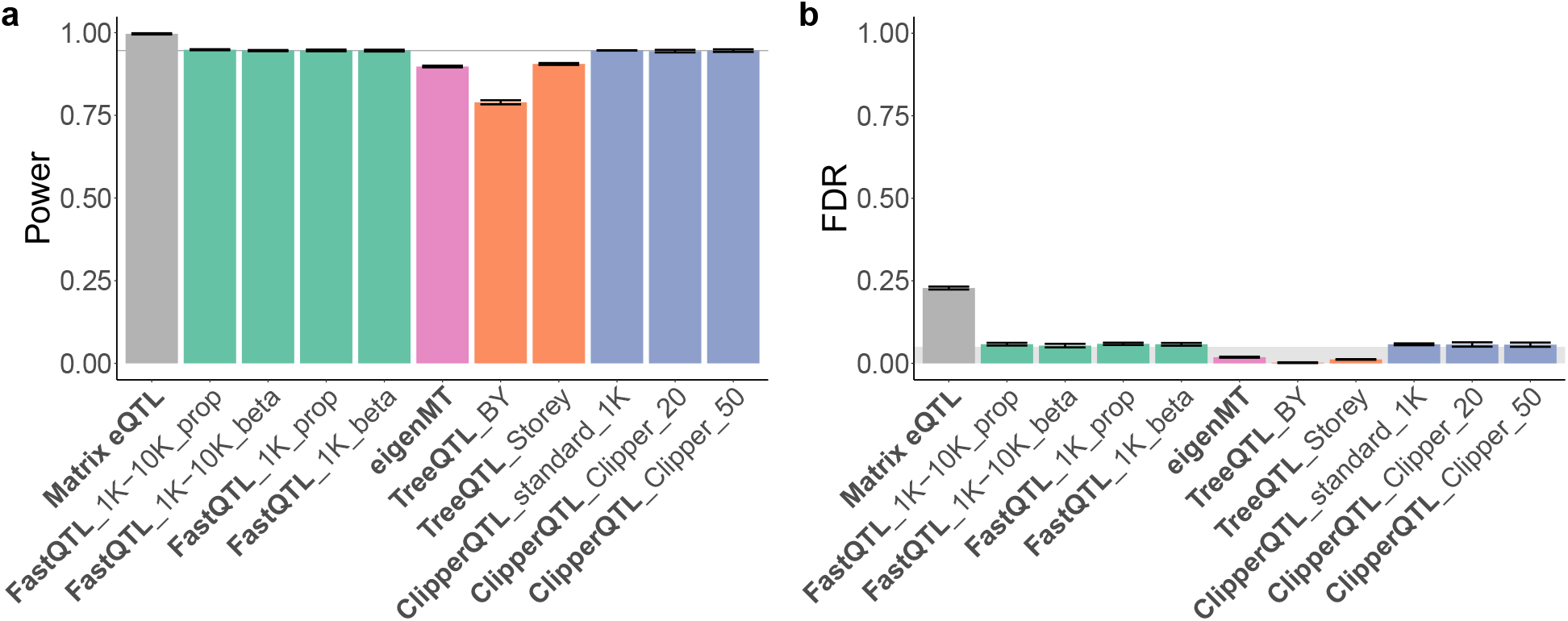
Power and FDR comparison of all 11 methods based on our simulation study (Table 1; Section 2.2). The target FDR is set at 0.05 (grey shaded area in **b**). The height of each bar represents the average across simulated data sets. Error bars indicate standard errors. In **a**, a horizontal line at the height of the bar for FastQTL_1K-10K beta is shown to help with visualization. All methods except Matrix eQTL can approximately control the FDR. FastQTL and ClipperQTL methods have higher power than eigenMT and TreeQTL methods.

In terms of run time comparison (Figure 2), we find that eigenMT has almost no computational advantage over FastQTL, and TreeQTL has no computational advantage over the standard variant of ClipperQTL (which is slower than the Clipper variant of ClipperQTL). Both the standard variant and the Clipper variant of ClipperQTL are orders of magnitude faster than FastQTL. In particular, the standard variant of ClipperQTL is about five times faster than FastQTL_1K_prop—the simplest FastQTL method—even though the algorithms are equivalent (Section 4.2); we attribute this to differences in software implementation. Compared to the default FastQTL method, the standard variant and the Clipper variant of ClipperQTL are about 50 times and 500 times faster, respectively.

### 2.2 Simulation results

In our simulation study, we roughly follow the data simulation in the second, more realistic simulation design of Zhou et al. [13], which roughly follows the data simulation in Wang et al. [17]. We simulate three data sets in total. Each data set is simulated according to Algorithm S5 with sample size *n* = 838, number of genes *p* = 1000, number of covariates 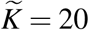, proportion of variance explained by genotype in eGenes `PVEGenotype` = 0.02, and proportion of variance explained by covariates `PVECovariates` = 0.5. All covariates are assumed to be known covariates.

The results from our simulation study are summarized in Figure 3. We confirm the finding in Huang et al. [16] that Matrix eQTL cannot control the FDR in the eGene identification problem. All other methods can approximately control the FDR. Further, FastQTL and ClipperQTL methods have higher power than eigenMT and TreeQTL methods, consistent with our real data results (Section 2.1).

## 3 Discussion

We have shown that ClipperQTL achieves a 500-fold or 50-fold increase in computational efficiency compared to FastQTL (depending on the variant used) without sacrificing power or precision. In contrast, other alternatives to FastQTL such as eigenMT and TreeQTL have lower power than FastQTL.

We propose two main variants of ClipperQTL: the standard variant and the Clipper variant. The standard variant is equivalent to FastQTL with the direct permutation scheme and proportions (Algorithm S1) and is suitable for a wide range of sample sizes. The Clipper variant uses the contrastive strategy developed in Clipper [11] (Algorithm 1) and is only recommended for data sets with large sample sizes (*>* 450).

Regarding which variant of ClipperQTL should be used when the sample size is large enough (*>* 450), we believe that if computational efficiency is a priority, then the Clipper variant should be used. However, if the study also contains smaller data sets, then the researcher may choose to use the standard variant on all data sets for consistency.

A possible extension of ClipperQTL lies in trans-eGene identification. Compared to cis-eGenes, trans-eGenes are currently identified in very small numbers [4], possibly due to the lack of power of existing approaches. Since the Clipper variant of ClipperQTL only needs 20 permutations for optimal performance and using only one permutation works almost as well (Section S3), there may be potential for ClipperQTL to be adapted for trans-eGene identification.

The R package ClipperQTL is available at https://github.com/heatherjzhou/ClipperQTL. Our work demonstrates the potential of the contrastive strategy developed in Clipper and provides a simpler and more efficient way of identifying cis-eGenes.

## 4 Methods

### 4.1 Problem

Here we describe the eGene identification problem and introduce the notations for this work.

The input data are as follows. Let *Y* denote the *n* × *p* fully processed gene expression matrix with *n* samples and *p* genes. For gene *j, j* = 1, *…, p*, the relevant genotype data is stored in *S* _*j*_, the *n* × *q* _*j*_ genotype matrix, where each column of *S* _*j*_ corresponds to a local common SNP for gene *j* (conceptually speaking; in reality, all genotype data may be stored in one file). Let *X* denote the *n* × *K* covariate matrix with *K* covariates. Using our analysis of GTEx’s Colon - Transverse expression data [4] (Section 2.1) as an example, we have *n* = 368, *p* = 25,379, *q* _*j*_ typically under 15,000, and *K* = 37, including eight known covariates and 29 inferred covariates (the number of inferred covariates is chosen via the BE algorithm [13, 14]; Section 2.1).

The assumption is that for *j* = 1, *…, p, Y* [, *j*], the *j*th column of *Y*, is a realization of the following random vector:

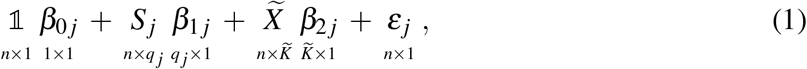

where 𝟙 denotes the *n* -*×* 1 matrix of ones, *S*_*j*_ is defined as above, 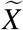 is the true covariate matrix (which *X* tries to capture), all entries of *β*_0 *j*_, *β*_1 *j*_, and *β*_2 *j*_ are fixed but unknown parameters, and *ε* _*j*_ is the random noise. In particular, it is assumed that at most a small number of entries of *β*_1 *j*_ are nonzero [17]. If all entries of *β*_1 *j*_ are zero, then gene *j is not* an eGene. On the other hand, if at least one entry of *β*_1 *j*_ is nonzero, then gene *j is* an eGene. The goal is to identify which of the *p* genes are eGenes given *Y*, 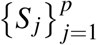, and *X* .

### 4.2 ClipperQTL

We propose two main variants of ClipperQTL: the standard variant and the Clipper variant. The standard variant is equivalent to FastQTL with the direct permutation scheme and proportions (Algorithm S1) and is suitable for a wide range of sample sizes. The Clipper variant uses the contrastive strategy developed in Clipper [11] (Algorithm 1) and is only recommended for data sets with large sample sizes (*>* 450). The development of ClipperQTL is discussed in Section S3. A key technical difference between the standard variant and the Clipper variant is that in the standard variant, gene expression is permuted first and then residualized, whereas in the Clipper variant, gene expression is residualized first and then permuted.

The main input parameter of ClipperQTL under both variants is *B*, the number of permutations. For the standard variant, *B* is set at 1000 by default. For the Clipper variant, we recommend setting *B* between 20 and 100 (Figures S3 and S4).

#### Algorithm 1

The Clipper variant of ClipperQTL

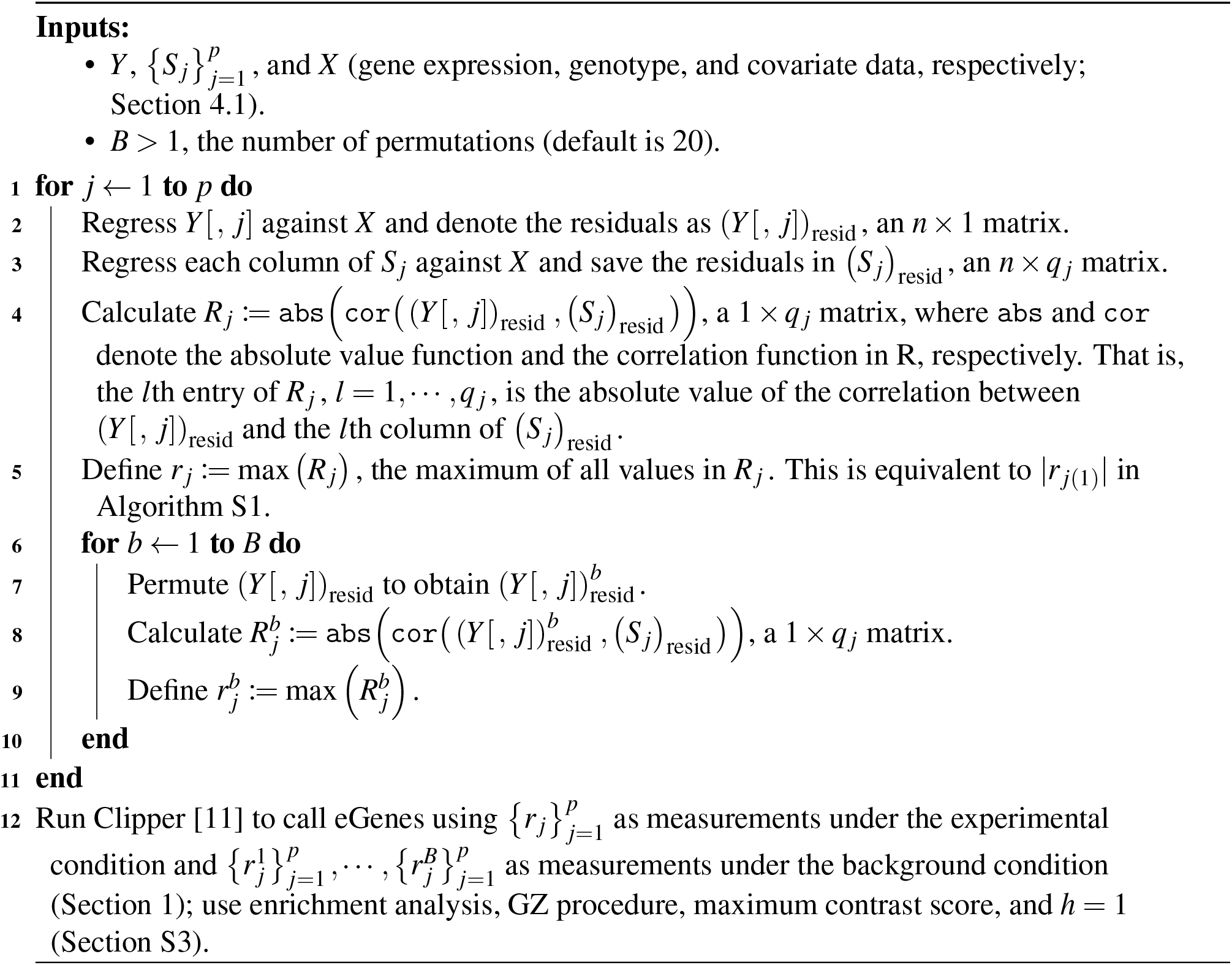

## Availability of data and materials

The R package ClipperQTL is available at https://github.com/heatherjzhou/ClipperQTL. The code used to generate the results in this work is available at https://doi.org/10.5281/zenodo.8259929. In addition, this work makes use of the following data and software:

- GTEx V8 public data [4], including fully processed gene expression matrices and known covariates, are downloaded from https://gtexportal.org/home/datasets.
- GTEx V8 protected data [4], specifically, the whole genome sequencing (WGS) phased genotype data, are downloaded from the AnVIL repository with an approved dbGaP application (see https://gtexportal.org/home/protectedDataAccess).
- FastQTL (https://github.com/francois-a/fastqtl, accessed October 29, 2020).
- Matrix eQTL R package Version 2.3 (https://cran.r-project.org/web/packages/MatrixEQTL, accessed March 6, 2023).
- eigenMT (https://github.com/joed3/eigenMT, accessed March 6, 2023).
- TreeQTL R package Version 2.0 (https://bioinformatics.org/treeqtl, accessed March 6, 2023).

## Authors’ contributions

H.J.Z, X.G., and J.J.L. conceived the project. H.J.Z. developed the method, performed the analyses and experiments, and wrote the software and manuscript. X.G. advised on the usage of Clipper. J.J.L. supervised the project. All authors participated in discussions and approved the final manuscript.

## Acknowledgments

The authors would like to thank former and current members of Junction of Statistics and Biology at UCLA for their valuable insight and suggestions.

## Declarations

### Ethics approval and consent to participate

Not applicable.

### Consent for publication

Not applicable.

## Supplementary materials for

### S1 Existing eGene identification methods

In this section, we review the existing eGene identification methods and describe the variants that we compare in this work (Table 1).

Recall the notations from Section 4.1. Let *Y* denote the *n × p* fully processed gene expression matrix with *n* samples and *p* genes. For gene *j, j* = 1, …, *p*, the relevant genotype data is stored in *S* _*j*_, the *n × q* _*j*_ genotype matrix, where each column of *S* _*j*_ corresponds to a local common SNP for gene *j* . Let *X* denote the *n×K* covariate matrix with *K* covariates.

#### S1.1 Matrix eQTL

Conceptually speaking, Matrix eQTL [15] works as follows: for *j* = 1,, *p* …, *l* = 1, …, *q* _*j*_, run the linear regression represented by the following R `lm()` formula:

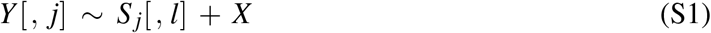

and obtain the *p*-value for the null hypothesis that the coefficient corresponding to *S* _*j*_[, *l*] is zero given the covariates; denote this *p*-value as *p* _*jl*_ . Therefore, a total of 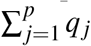 *p*-values are obtained, one for each gene-SNP pair. Matrix eQTL then uses the Benjamini-Hochberg (BH) procedure [18] on these *p*-values to call significant gene-SNP pairs [15]. To call eGenes using Matrix eQTL in this work, we call as eGenes all genes that appear at least once in the significant gene-SNP pairs.

In reality, Matrix eQTL uses the following equivalent approach to obtain the *p*-values, which is more computationally efficient due to the overlap of local common SNPs across genes. For *j* = 1, *…, p, l* = 1, *…, q* _*j*_, first, regress the gene expression against the covariates:

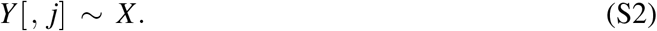

Second, regress the genotype against the covariates:

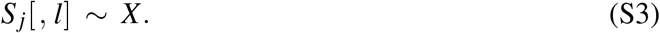

Then, calculate the Pearson correlation between the expression residuals from (S2) and the genotype residuals from (S3) and denote it as *r* _*jl*_ . This is the partial correlation between *Y* [, *j*] and *S* _*j*_[, *l*] conditional on *X* .

Lastly, convert the partial correlation to a test statistic using

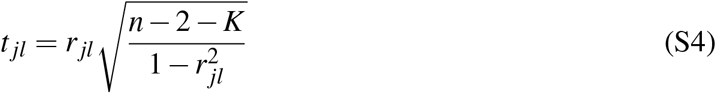

and convert the test statistic to a *p*-value using

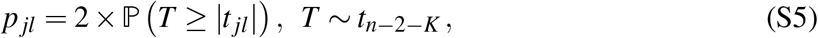

where ℙ denotes probability, |*t* _*jl*_| denotes the absolute value of *t* _*jl*_, and *T* is a random variable following the *t*-distribution with *n*− 2 −*K* degrees of freedom.

Notably, the larger |*r* _*jl*_| (the absolute value of *r* _*jl*_), the smaller *p* _*jl*_, and vice versa. Both FastQTL (the variants using proportions; Section S1.2) and ClipperQTL (the standard variant; Section 4.2) make use of this fact.

#### S1.2 FastQTL

There are four main ways to use FastQTL [6], depending on (1) whether the direct or the adaptive permutation scheme is used and (2) whether proportions or beta approximation is used. The direct permutation scheme with either proportions or beta approximation is summarized in Algorithm S1. The adaptive permutation scheme is identical except the number of permutations is chosen adaptively between *B*_min_ and *B*_max_ (two input parameters) for each gene rather than directly inputted (see Ongen et al. [6] for details).

The default way of using FastQTL is to use the adaptive permutation scheme (*B*_min_ = 1000 and *B*_max_ = 10,000) with beta approximation [4, 6]. In total, we compare four ways of using FastQTL in this work including the default approach: FastQTL_1K-10K_prop, FastQTL_1K-10K beta (the default), FastQTL_1K_prop, and FastQTL_1K_beta (see Table 1). That is, the number of permutations is either fixed at 1000 or chosen adaptively between 1000 and 10,000 for each gene, and either proportions or beta approximation is used.

In addition to identifying eGenes, FastQTL can also output significant gene-SNP pairs. We summarize the algorithm for this in Algorithm S2.

##### Algorithm S1

The direct permutation scheme of FastQTL

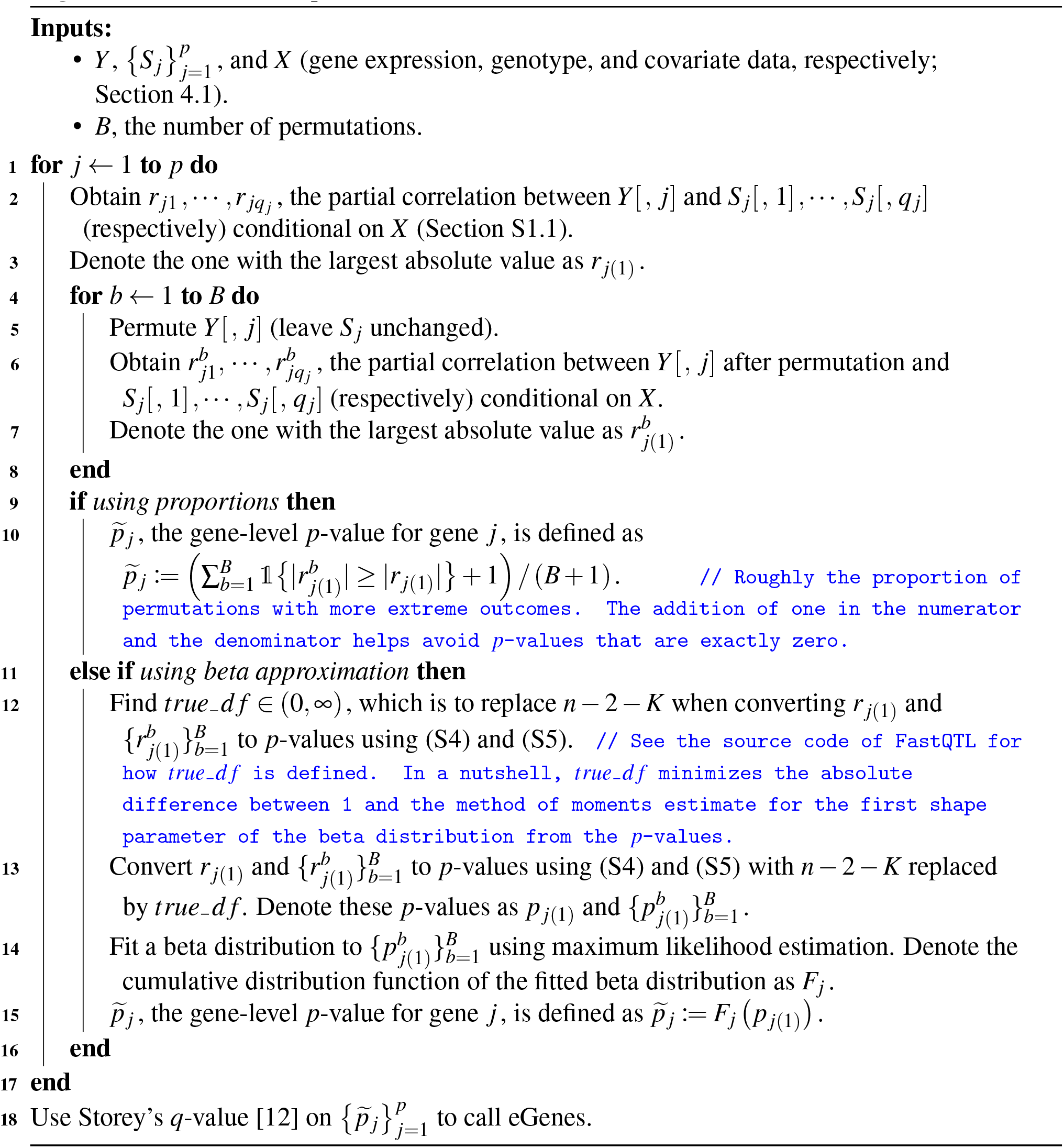

##### Algorithm S2

Identification of significant gene-SNP pairs in FastQTL

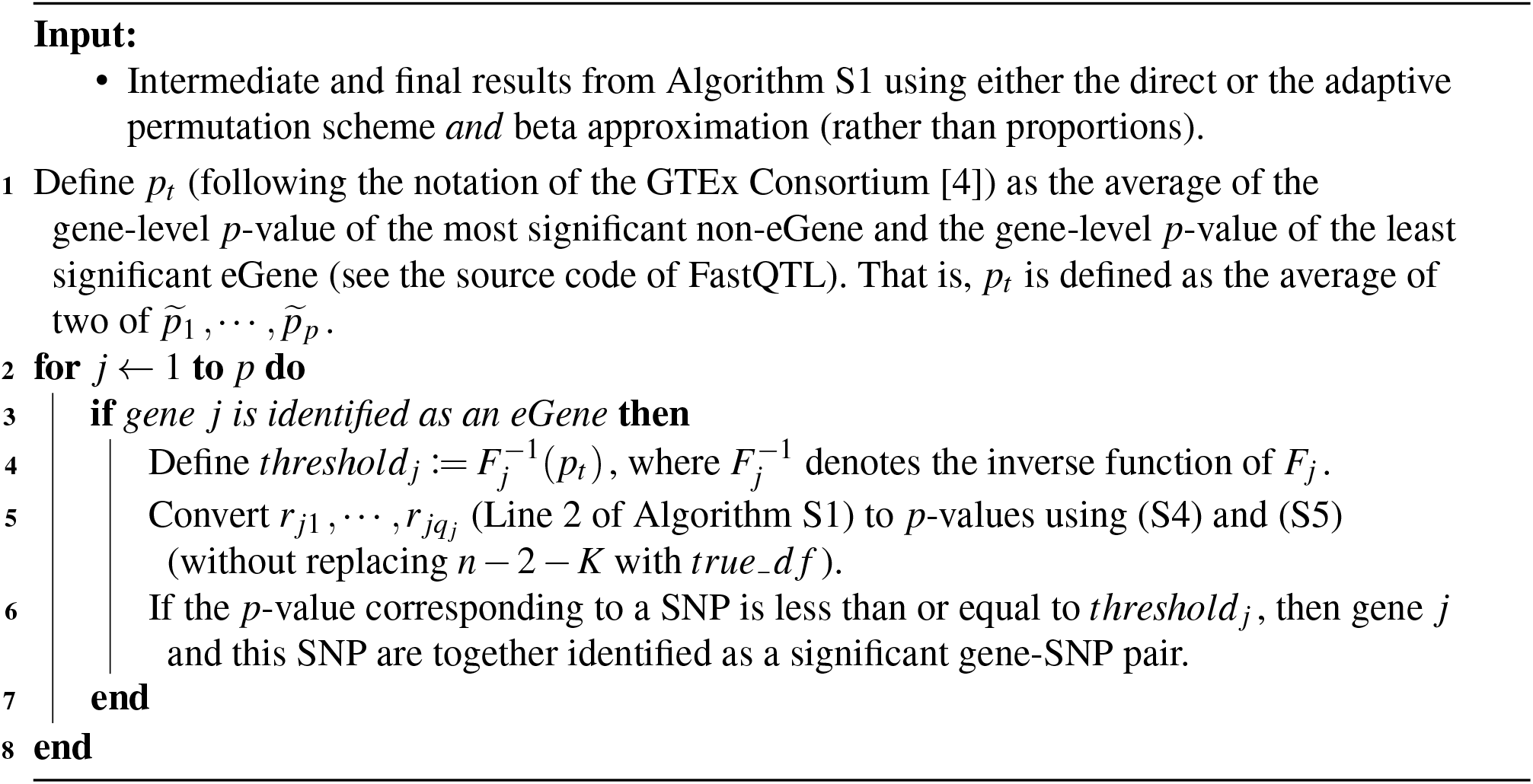

**Figure S1:**
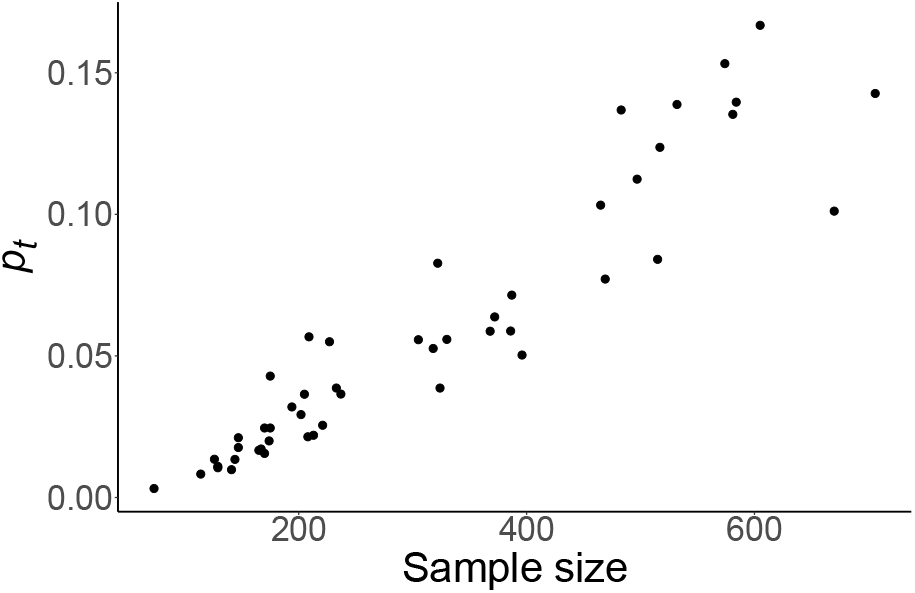
Scatter plot of *p*_*t*_ (Algorithm S2) from FastQTL_1K-10K beta versus sample size in GTEx expression data [4] (see Section 2.1 for the analysis details). This scatter plot contains 49 dots, each corresponding to a tissue. We see that *p*_*t*_ increases roughly linearly with sample size.

#### S1.3 eigenMT

We summarize eigenMT [7] in Algorithm S3. After obtaining the gene-level *p*-values, eigenMT does not specify what method to use to control the false discovery rate when calling eGenes. Therefore, in this work, we use Storey’s *q*-value [12] following FastQTL (Section S1.2). In addition to producing the gene-level *p*-values, eigenMT also outputs the most significantly associated SNP for each gene.

##### Algorithm S3

eigenMT

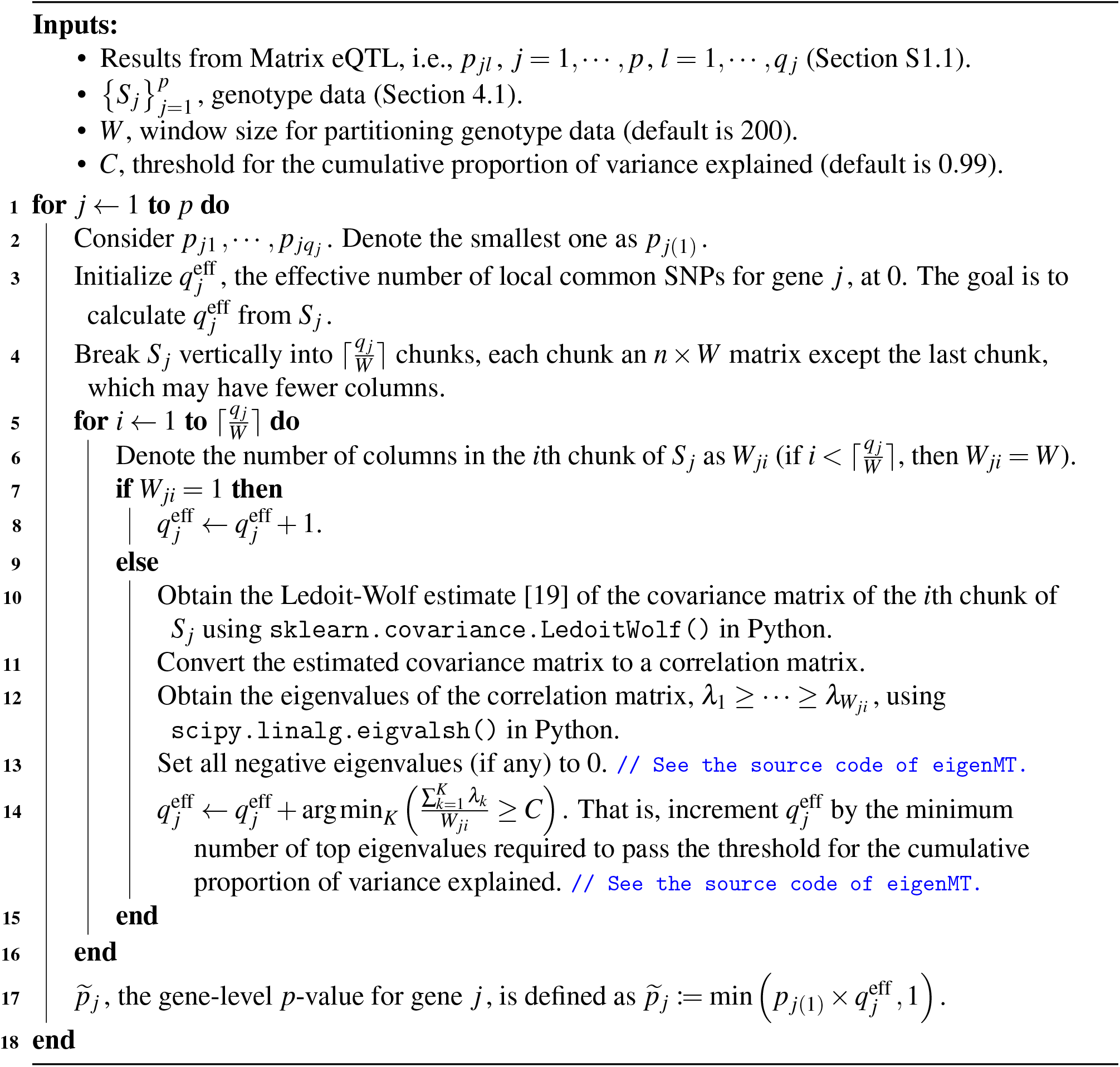

#### S1.4 TreeQTL

TreeQTL [8] uses Simes’ rule [10] to calculate a gene-level *p*-value for each gene (Algorithm S4). After obtaining the gene-level *p*-values, TreeQTL allows the user to use Bonferroni correction, BH [18], or Benjamini-Yekutieli (BY) [20] to call eGenes (the default is BY).

We compare two variants of TreeQTL in this work: TreeQTL BY (the default) and TreeQTL_Storey (see Table 1). In TreeQTL_Storey, we use Storey’s *q*-value [12] on the gene-level *p*-values to call eGenes, following FastQTL (Section S1.2). We do not include variants of TreeQTL using Bonferroni correction or BH in our comparison because Bonferroni correction aims to control the family-wise error rate rather than the false discovery rate, and BH is more stringent than Storey’s *q*-value (we show that even TreeQTL_Storey has lower power than FastQTL; Figures 1 and 3).

##### Algorithm S4

TreeQTL

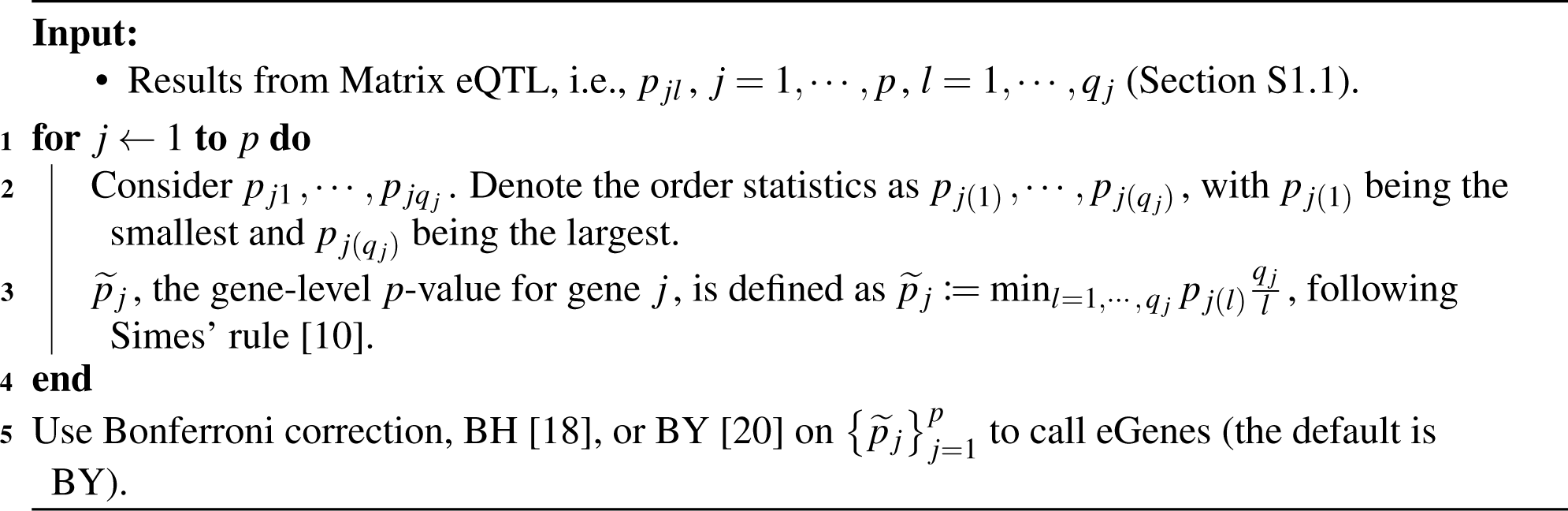

### S2 Data simulation

In our simulation study, we roughly follow the data simulation in the second, more realistic simulation design of Zhou et al. [13], which roughly follows the data simulation in Wang et al. [17]. We simulate three data sets in total. Each data set is simulated according to Algorithm S5 with the following attributes:

- Sample size, *n* = 838.
- Number of genes, *p* = 1000.
- Number of covariates, 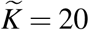.
- Proportion of variance explained by genotype in eGenes, `PVEGenotype` = 0.02.
- Proportion of variance explained by covariates, `PVECovariates` = 0.5.

**Table S1:** In our data simulation (Algorithm S5), the number of effect SNPs [13, 17] for each gene is sampled based on this probability table (the second column sums to one). This table is summarized from GTEx’s independent cis-eQTL analysis [4] (see Figure S2 of Zhou et al. [13]). A gene is an eGene if and only if its number of effect SNPs is greater than zero.

| # of effect SNPs | Probability |
| --- | --- |
| 0 | 0.35483532 |
| 1 | 0.34962617 |
| 2 | 0.18326554 |
| 3 | 0.07072812 |
| 4 | 0.02498728 |
| 5 | 0.01655758 |

#### Algorithm S5

Simulation of one data set

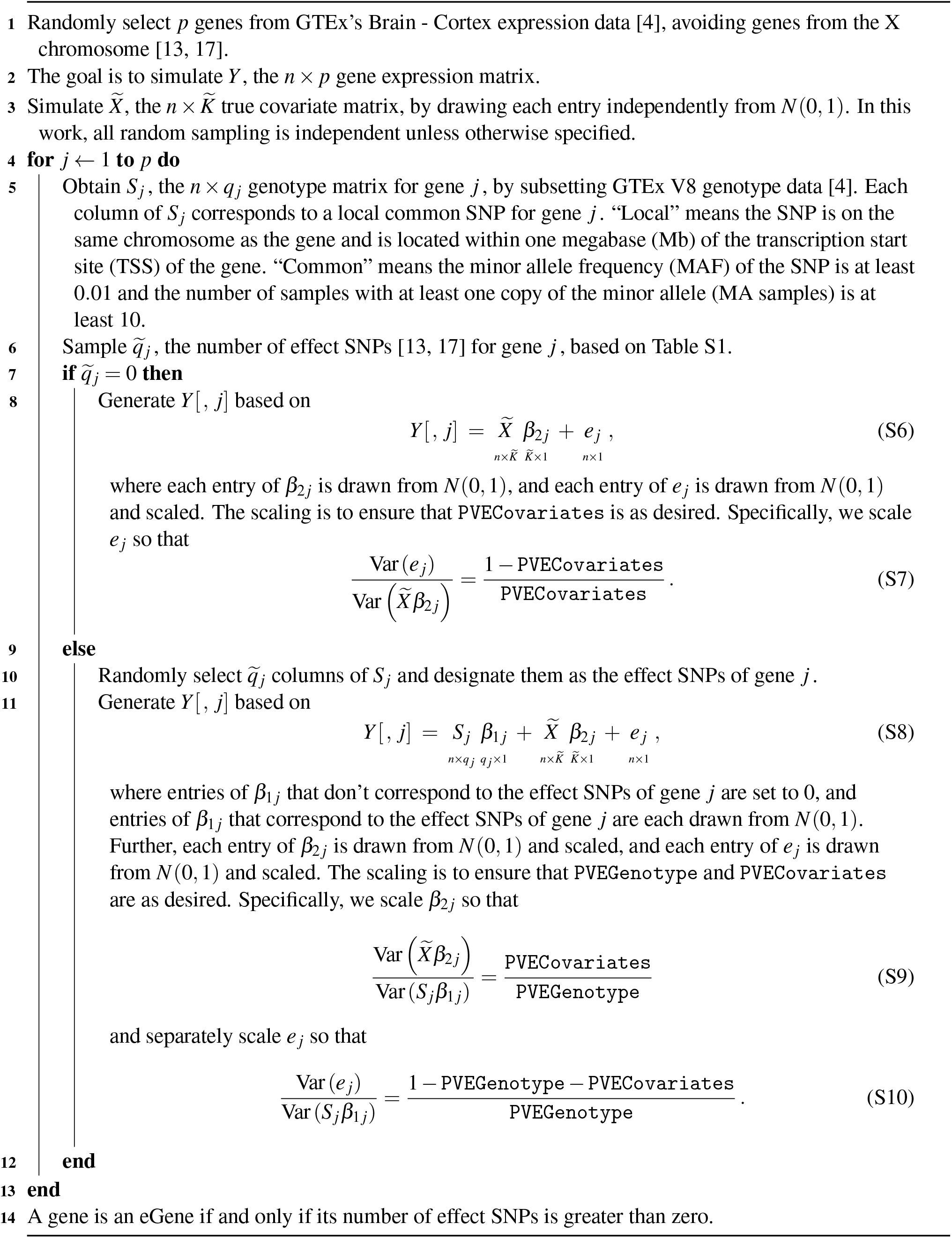

### S3 Development of ClipperQTL

Here we describe the development of ClipperQTL.

Clipper [11] has four main technical parameters:

- Analysis: enrichment vs. differential analysis.
- Procedure: Barber-Cande’s (BC) vs. Gimenez-Zou (GZ) procedure.
- Contrast score: maximum vs. difference (i.e., minus) contrast score.
- *h* (only applicable under the GZ procedure, not the BC procedure).

In addition, in ClipperQTL, we can control *B*, the number of permutations (ClipperQTL terminology), i.e., the number replicates under the background condition (Clipper terminology).

In ClipperQTL, we use enrichment analysis rather than differential analysis because in identifying eGenes, the alternative hypothesis is that the expectation of the maximum absolute correlation from the original expression data is *greater than* (rather than merely different from) the expectation of the maximum absolute correlation from permuted expression data.

In ClipperQTL, the number of replicates under the experimental condition is fixed at one because we only have one set of the original, unpermuted expression data. Therefore, if *B* = 1, then we only need to consider the BC procedure (in enrichment analysis, if the number of replicates under the experimental condition and the number of replicates under the background condition are both one, then the GZ procedure with either maximum or difference contrast score reduces to the BC procedure with difference contrast score); if *B >* 1, then we only need to consider the GZ procedure (in enrichment analysis, the BC procedure is only applicable when the number of replicates under the experimental condition and the number of replicates under the background condition are equal [11]). In other words, *B* determines the procedure we need to consider.

Therefore, we explore different combinations of *B*, contrast score, and *h* (only applicable under the GZ procedure). We find that for data sets with small sample sizes (*<* 450), no combination works well consistently, but for data sets with large sample sizes (*>* 450), *B* between 20 and 100, maximum contrast score, and *h* = 1 works well (*B* = 1 and maximum contrast score works almost as well; details not shown). Therefore, the Clipper variant of ClipperQTL is only recommended for data sets with large sample sizes (*>* 450; Section 4.2). It always uses enrichment analysis, GZ procedure, maximum contrast score, and *h* = 1 (Algorithm 1), and the user is recommended to set *B* between 20 and 100 (Section 4.2).

**Figure S2:**
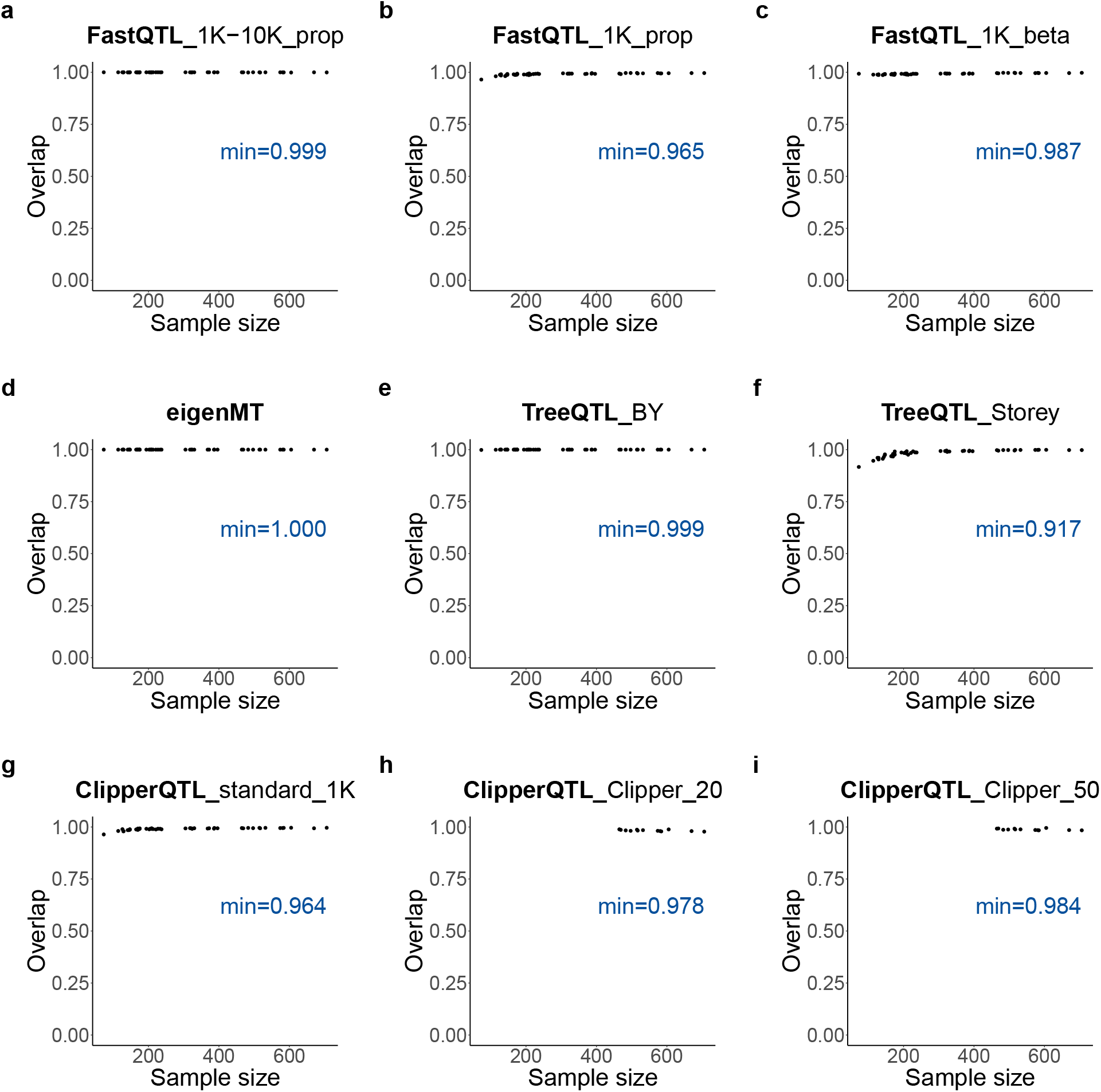
Overlap between eGenes identified by various methods and eGenes identified by FastQTL_1K-10K beta—the default FastQTL method—in GTEx expression data [4] (Table 1; see Section 2.1 for the analysis details). Each dot corresponds to a tissue. Given two sets, *A* and *B*, the overlap is defined as |*A* ∩ *B*| */* min(|*A*|, |*B*|), where | · | denotes the cardinality of a set. That is, the overlap between two sets is defined as the size of the intersection divided by the size of the smaller set. **b, c, g** The overlap is slightly lower when the sample size is smaller. This can be explained by the fact that power is generally lower when the sample size is smaller [4]. **h, i** Only tissues with sample sizes ≥ 465 are shown (Figure 1).

**Figure S3:**
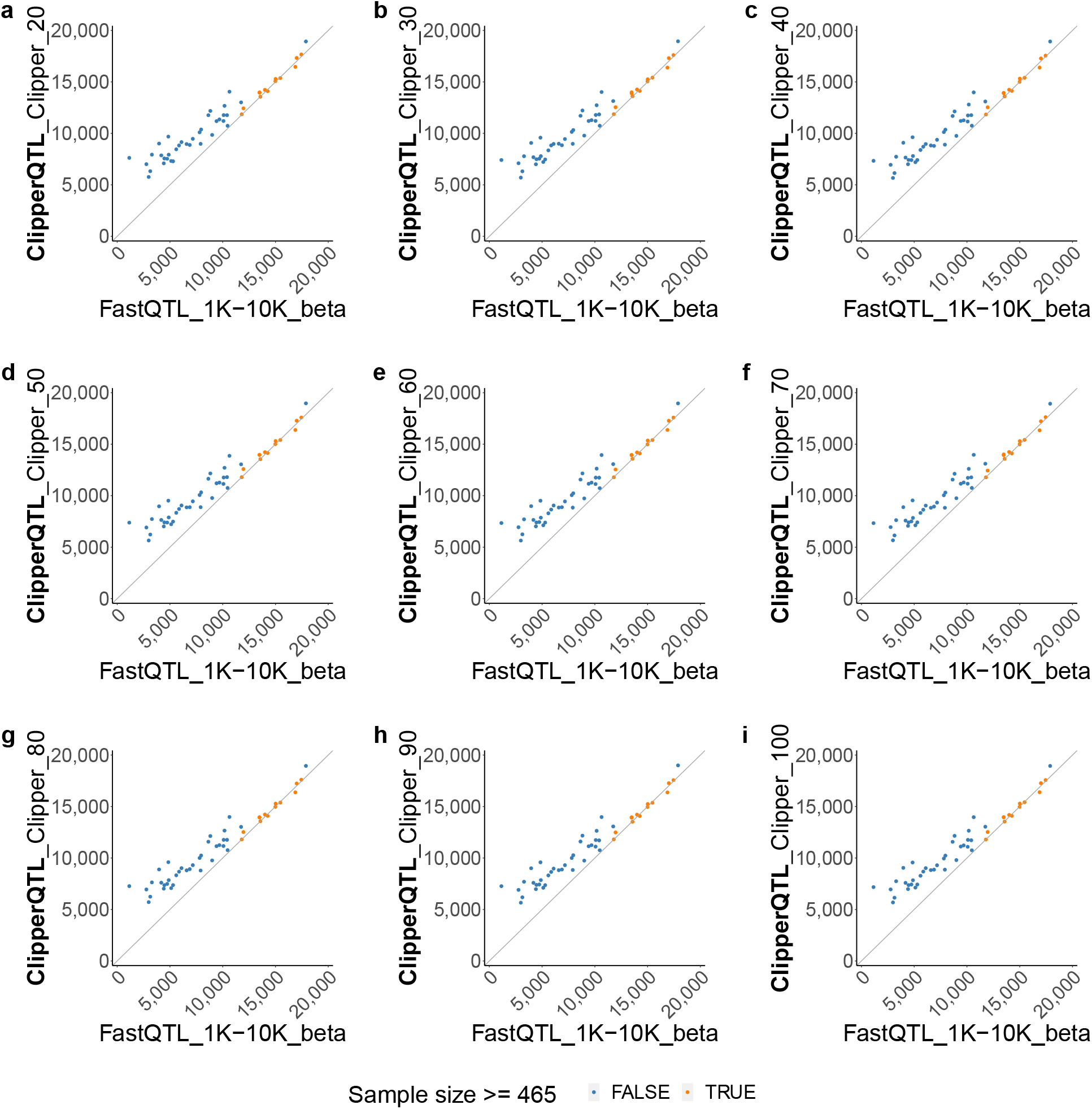
Number of eGenes comparison between ClipperQTL_Clipper with different *B*’s and the default FastQTL method based on GTEx expression data [4] (the analysis details are as described in Section 2.1). Each dot corresponds to a tissue. The *x*-axis and *y*-axis both represent numbers of eGenes identified by different methods. Diagonal lines through the origin are shown to help with visualization. ClipperQTL_Clipper with *B* between 20 and 100 works well for tissues with large sample sizes (Section 4.2). We use 465 as the sample size cutoff because the next largest sample size is 396.

**Figure S4:**
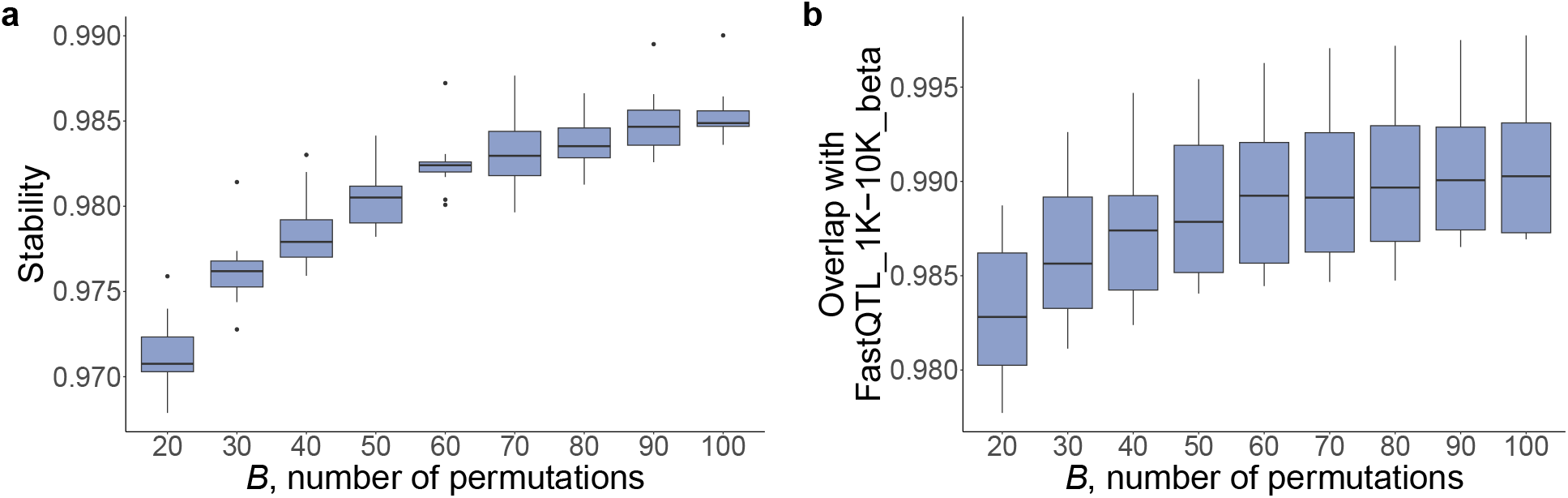
Comparison of ClipperQTL_Clipper with different *B*’s. Each box plot contains 13 data points, corresponding to the 13 tissues in GTEx expression data [4] with sample sizes ≥ 465 (the analysis details are as described in Section 2.1). As *B* increases from 20 to 100, the stability of ClipperQTL_Clipper (**a**) and the discovery overlap with the default FastQTL method (**b**) both increase slightly. See Figure S2 for our definition of overlap. Stability is a measure of how much the result of a method depends on the random seed; the higher the stability, the less the result varies with respect to the random seed. Specifically, to calculate the stability of a method (e.g., ClipperQTL_Clipper_20), we run the method 10 times with 10 different seeds. We divide the 10 runs into 5 pairs. For each pair, we calculate the overlap between the two sets of identified eGenes. The stability of the method is calculated as the average of the 5 overlaps.

## References

[1] Eddie Cano-Gamez and Gosia Trynka. From GWAS to function: Using functional genomics to identify the mechanisms underlying complex diseases. Frontiers in Genetics, 11:424, 2020.

[2] Youqiong Ye, Zhao Zhang, Yaoming Liu, Lixia Diao, and Leng Han. A multi-omics perspective of quantitative trait loci in precision medicine. Trends in Genetics, 36(5):318–336, 2020.

[3] GTEx Consortium. Genetic effects on gene expression across human tissues. Nature, 550 (7675):204–213, 2017.

[4] GTEx Consortium. The GTEx Consortium atlas of genetic regulatory effects across human tissues. Science, 369(6509):1318–1330, 2020.

[5] Lei Li, Kai-Lieh Huang, Yipeng Gao, Ya Cui, Gao Wang, Nathan D. Elrod, Yumei Li, Yiling Elaine Chen, Ping Ji, Fanglue Peng, William K. Russell, Eric J. Wagner, and Wei Li. An atlas of alternative polyadenylation quantitative trait loci contributing to complex trait and disease heritability. Nature Genetics, 53(7):994–1005, 2021.

[6] Halit Ongen, Alfonso Buil, Andrew Anand Brown, Emmanouil T. Dermitzakis, and Olivier Delaneau. Fast and efficient QTL mapper for thousands of molecular phenotypes. Bioinformatics, 32(10):1479–1485, 2016.

[7] Joe R. Davis, Laure Fresard, David A. Knowles, Mauro Pala, Carlos D. Bustamante, Alexis Battle, and Stephen B. Montgomery. An efficient multiple-testing adjustment for eQTL studies that accounts for linkage disequilibrium between variants. The American Journal of Human Genetics, 98(1):216–224, 2016.

[8] C. B. Peterson, M. Bogomolov, Y. Benjamini, and C. Sabatti. TreeQTL: Hierarchical error control for eQTL findings. Bioinformatics, 32(16):2556–2558, 2016.

[9] Amaro Taylor-Weiner, FranÇois Aguet, Nicholas J. Haradhvala, Sager Gosai, Shankara Anand, Jaegil Kim, Kristin Ardlie, Eliezer M. Van Allen, and Gad Getz. Scaling computational genomics to millions of individuals with GPUs. Genome Biology, 20(1):228, 2019.

[10] R. J. Simes. An improved Bonferroni procedure for multiple tests of significance. Biometrika, 73(3):751–754, 1986.

[11] Xinzhou Ge, Yiling Elaine Chen, Dongyuan Song, MeiLu McDermott, Kyla Woyshner, Antigoni Manousopoulou, Ning Wang, Wei Li, Leo D. Wang, and Jingyi Jessica Li. Clipper: P-value-free FDR control on high-throughput data from two conditions. Genome Biology, 22 (1):288, 2021.

[12] John D. Storey and Robert Tibshirani. Statistical significance for genomewide studies. Proceedings of the National Academy of Sciences, 100(16):9440–9445, 2003.

[13] Heather J. Zhou, Lei Li, Yumei Li, Wei Li, and Jingyi Jessica Li. PCA outperforms popular hidden variable inference methods for molecular QTL mapping. Genome Biology, 23(1):210, 2022. 10.1186/s13059-022-02761-4.

[14] Andreas Buja and Nermin Eyuboglu. Remarks on parallel analysis. Multivariate Behavioral Research, 27(4):509–540, 1992.

[15] Andrey A. Shabalin. Matrix eQTL: Ultra fast eQTL analysis via large matrix operations. Bioinformatics, 28(10):1353–1358, 2012.

[16] Qin Qin Huang, Scott C Ritchie, Marta Brozynska, and Michael Inouye. Power, false discovery rate and Winner’s Curse in eQTL studies. Nucleic Acids Research, 46(22):e133–e133, 2018.

[17] Gao Wang, Abhishek Sarkar, Peter Carbonetto, and Matthew Stephens. A simple new approach to variable selection in regression, with application to genetic fine mapping. Journal of the Royal Statistical Society: Series B (Statistical Methodology), 82(5):1273–1300, 2020.

[18] Yoav Benjamini and Yosef Hochberg. Controlling the false discovery rate: A practical and powerful approach to multiple testing. Journal of the Royal Statistical Society: Series B (Methodological), 57(1):289–300, 1995.

[19] Olivier Ledoit and Michael Wolf. A well-conditioned estimator for large-dimensional covariance matrices. Journal of Multivariate Analysis, 88(2):365–411, 2004.

[20] Yoav Benjamini and Daniel Yekutieli. The control of the false discovery rate in multiple testing under dependency. The Annals of Statistics, 29(4):1165–1188, 2001.

